# Double nicking by RNA-directed Cascade-nCas3 for high-efficiency large-scale genome engineering

**DOI:** 10.1101/2021.07.12.451994

**Authors:** Yile Hao, Qinhua Wang, Jie Li, Shihui Yang, Lixin Ma, Yanli Zheng, Wenfang Peng

## Abstract

New CRISPR-based genome editing technologies are developed to continuedly drive advances in life sciences, which, however, are predominantly derived from systems of Type II CRISPR-Cas9 and Type V CRISPR-Cas12a for eukaryotes. Here we report a novel CRISPR-n(nickase)Cas3 genome editing tool established upon an endogenous Type I system of *Zymomonas mobilis*. We demonstrate that nCas3 variants can be created by alanine-substituting any catalytic residue of the Cas3 helicase domain. While nCas3 overproduction via plasmid shows severe cytotoxicity; an *in situ* nCas3 introduces targeted double-strand breaks, facilitating genome editing, without visible cell killing. By harnessing this CRISPR-nCas3, deletion of genes or genomic DNA stretches can be consistently accomplished with near-100% efficiencies, including simultaneous removal of two large genomic fragments. Our work describes the first establishment of a CRISPR-nCas3-based genome editing technology, thereby offering a simple, easy, yet useful approach to convert many endogenous Type I systems into advanced genome editing tools. We envision that many CRISPR-nCas3-based toolkits would be soon available for various industrially important non-model bacteria that carry active Type I systems to facilitate high-throughput prokaryotic engineering.

---

CRISPR-Cas systems are a group of RNA-guided machineries that defend their prokaryotic hosts against invasive genetic elements in a programmable manner (Barrangou et al. 2007; Brouns et al. 2008). The targetable DNA-binding Cas nucleases are therein applied in generating double-stranded DNA breaks (DSBs) at specific chromosomal loci, stimulating the host repair mechanisms, including homology-directed repair (HDR) and non-homologous end joining (NHEJ), to bring about designed or error-prone genomic alterations (Anzalone et al. 2020). Such applications have been currently focused on the compact Class 2 systems with a single Cas effector on account of their simplicity and hence ease of heterologous use (Makarova et al. 2020). Among Class 2 systems, the notable CRISPR-Cas9 from *Streptococcus pyogenes* has been pioneered successful genome editing in various organisms or cell lines (Graham and Root 2015; Barrangou and Doudna 2016). The success of wild-type Cas9-based applications has also fuelled the development of the technologies based on its derivatives, such as the nCas9 (Cas9 nickase) that possesses several advantages over the original (Anzalone et al. 2020). For instance, a paired-nCas9 strategy can be used to greatly enhance DNA targeting specificity and consequently lower off-targeting in genome editing (Ran et al. 2013). Additionally, nCas9 can help deaminases to yield more predictable and precise genome editing compared with wild-type Cas9-based editing (Nishida et al. 2016).

Despite of the versatility and robustness of the CRISPR-Cas9/nCas9 technologies, their applications in prokaryotes have been rather limited, because overexpressing the exogenous Cas proteins in most bacteria could be cytotoxic and would lead to failure in yielding colonies (Vento et al. 2019). As an alternative strategy, several Type I CRISPR-Cas3 systems belonging to Class 1 have been exploited to work as “built-in” genome editing tools in their native hosts (Zheng et al. 2020; Xu et al. 2021), including Type I-A of *Sulfolobus islandicus* (Li et al. 2016), Type I-B of *Haloarcula hispanica* (Cheng et al. 2017) and *Clostridium* species (Pyne et al. 2016; Zhang et al. 2018), Type I-C of *Pectobacterium aeruginosa* (Csorgo et al. 2020), Type I-E of *Streptococcus thermophilus* (Canez et al. 2019) and *Lactobacillus crispatus* (Hidalgo-Cantabrana et al. 2019), and Type I-F of *Pectobacterium* species (Vercoe et al. 2013; Xu et al. 2019), and *Zymomonas mobilis* (Zheng et al. 2019), where the processive Cas3 nuclease-helicase was used to generate chromosomal injuries. Recent studies have also employed Type I-D and I-E systems for DNA cleavage in plants (Osakabe et al. 2020) and human cells (Cameron et al. 2019; Dolan et al. 2019; Morisaka et al. 2019), respectively, and Type I-E and I-F systems for gene expression modulation in human cells (Pickar-Oliver et al. 2019; Chen et al. 2020), further broadening the applicability of CRISPR-Cas3-based technologies. These accomplishments have paved a new possibility to develop advanced CRISPR-nCas3 toolkits based on endogenous Type I systems. Yet, to the best of our knowledge, no CRISPR-nCas3-based technology has been currently available.

We have previously accomplished genome engineering with the endogenous Type I-F CRISPR-Cas3 system of *Z. mobilis* ZM4. In the work, the editing options concerning single genes, including knockout, replacement, and *in situ* nucleotide substitutions, yielded 100% efficiencies; whereas others did not, for example, while at most 50% efficiency could be got in the deletion of a large genomic fragment (ca. 5‰ of the genome) (Zheng et al. 2019). Here we have, for the first time, developed a CRISPR-nCas3 genome editing tool, which has enabled large-scale genomic deletions with near-100% efficiencies that is currently hardly achievable using other methodologies. In addition, this tool has allowed for simultaneous deletion of two large genomic fragments with an efficiency of up to 75%, showing its great potential to sever as a versatile tool for high-throughput metabolic engineering proctices.

## Results

### Inactivation of the helicase domain converts the Cas3 nuclease-helicase into a nickase

Cas3 possesses activities of ssDNA-specific nuclease and ATP-dependent helicase, being responsible for target cleavage and degradation in Type I CRISPR-Cas systems (He et al. 2020). The nuclease domain of Cas3 initially nicks the target sequence within the ssDNA region of an R-loop generated upon Cascade-binding and crRNA invasion. Subsequently, by consuming ATP, Cas3 unwinds the dsDNA starting at the nicked site via its helicase domain to further provide ssDNA substrate for its nuclease domain, eventually leading to complete target degradation (Sinkunas et al. 2011; Hidalgo-Cantabrana and Barrangou 2020). We reasoned that mutating the catalytic residues of the helicase domain might convert Cas3 into a nickase (nCas3), which could no longer unwind the dsDNA due to the loss of its ATPase activity. To verify this assumption, we opted to create nCas3 variants and assess their capability on plasmid DNA nicking.

Amino acid sequence alignment of the Cas3 from *Z. mobilis* (*Zmo*Cas3), actually a Cas2-Cas3 fusion encoding by the *cas2/3* gene (Zheng et al. 2019), with several reported Cas3 homologs had revealed its characteristic helicase motifs (I, II and VI) coordinating ATP binding and hydrolysis (Sinkunas et al. 2011; Gong et al. 2014) (**Fig. 1a** and **Figure S1**). We therefore designed alanine substitution of conserved residues including K458 located in motif I, D608 in motif II, and R887 in motif VI (**Fig. 1b**). The variants, as well as the wild-type *Zmo*Cas3, could be recombinantly produced in *Escherichia coli* as soluble proteins (**Fig. 1c**), and each of which, together with the Cascade-crRNA complex, was incubated with a 3,283-bp negatively supercoiled (NS) plasmid, pL2R (Zheng et al. 2019) (**Table S1**), bearing a functional 5’-CCC-3’ PAM-preceded protospacer sequence. The treated DNAs were subsequently subjected to electrophoreses using agarose gels. As shown in **Fig. 1d**, following nicking the NS plasmid into an open circle (OC) DNA, the wild-type *Zmo*Cas3 (wt) eventually degraded the plasmid DNA completely; whereas the nCas3 variants gradually nicked the NS plasmid DNA into the OC version. Linear (L) DNAs were also observed, indicative of the occurrence of DSBs. Possibly, in the finite *in vitro* reactions the nuclease domain of free nCas3 variants could have occasionally touched and cut the opposite strand of the nicked site. These results suggested that all these variants are nCas3s.

**Fig. 1.**
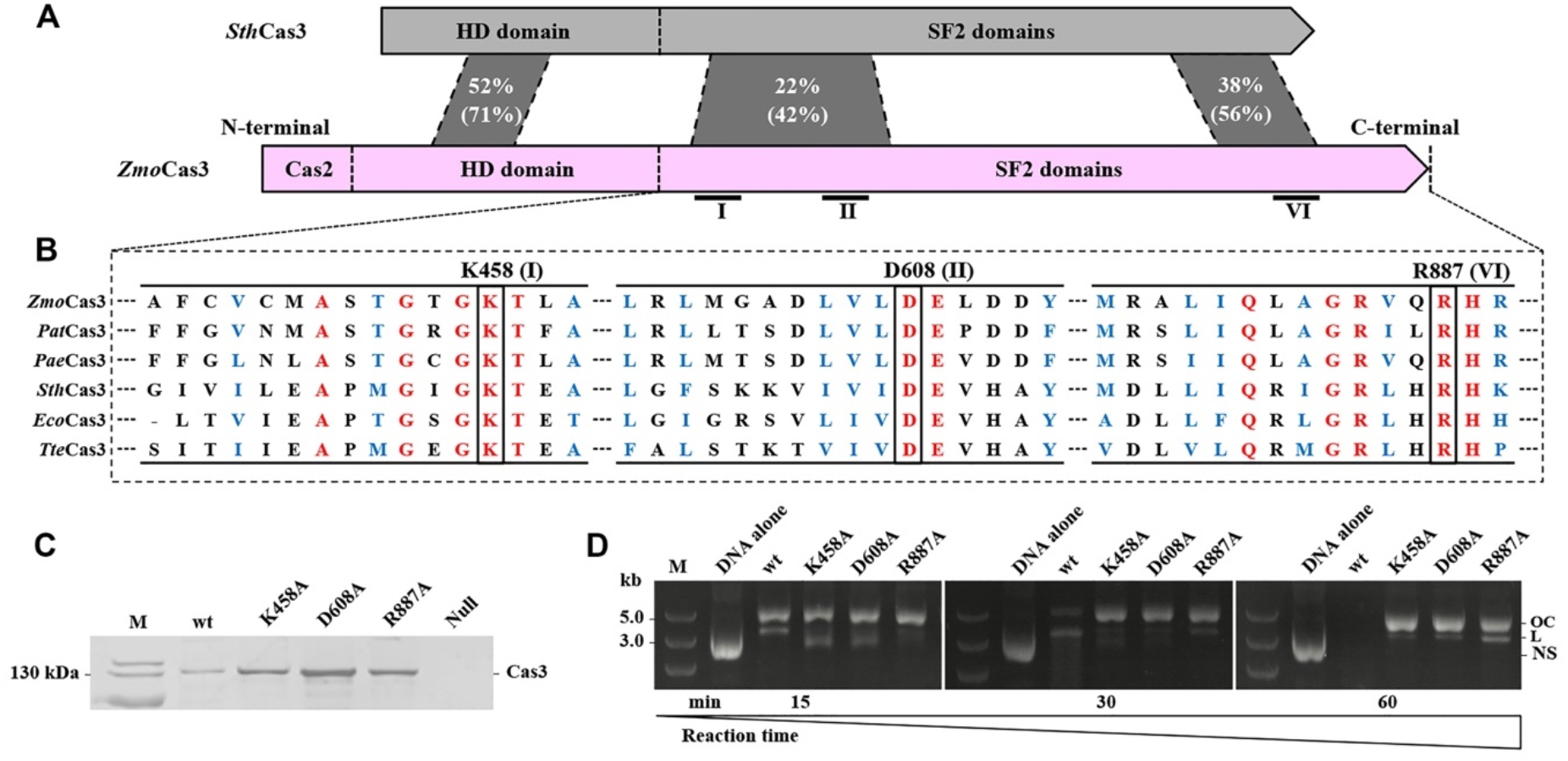
Construction of Cas3 nickase mutants. (**A**) Schematic organization of Cas3 proteins from *Zymomonas mobilis* ZM4 (*Zmo*Cas3) and *Streptococcus thermophilus* DGCC7710 (*Sth*Cas3). Domain architecture of the Cas3 proteins identified by *in silico* analysis is shown as pink (*Zmo*Cas3) and grey (*Sth*Cas3) boxes, respectively. Percentage of identical and similar (in parenthesis) residues between conserved sequence blocks is shown. For *Zmo*Cas3, Cas2 denotes the N-terminally fused Cas2 domain; HD domain denotes HD-type phosphohydrolase/nuclease domain; SF2 domains denote helicase domains. (**B**) Locations of the conserved helicase motifs are indicated (I, II, and VI) which were identified by alignment of Cas3 proteins from different CRISPR-Cas systems of Type I-E and I-F. Conserved residues characteristic of each motif (K458 of motif I, D608 of motif II, and R887 of motif VI, respectively) being subjected to alanine mutagenesis are indicated above the corresponding positions. *Pat*, *Pectobacterium atrosepticum*; *Pae*, *Pseudomonas aeruginosa*; *Eco*, *Escherichia coli* K-12; *Tte*, *Thermobaculum terrenum*. (**C**) Coomassie blue-stained SDS-PAGE of purified Cas3 proteins expressed in *E. coli*, including the wild-type *Zmo*Cas3 (wt) and three Cas3 nickase candidates. Null, *E. coli* BL21 (DE3) cells carrying the cloning vector pET28a; M, protein size marker. (**D**) Analyses of plasmid DNA cleavage by the purified Cas3 proteins as indicated in (**C**) via electrophoreses using agarose gels. OC, open circle; L, linear; NS, negatively supercoiled; M, DNA size marker.

### Overexpression of nCas3 has potent killing effect on *Z. mobilis* cells

Having determined the nickase nature of the nCas3 mutants, we next studied whether they could be employed to make DSBs through double nicking for genome editing in *Z. mobilis*. We chose the *ZMO0038* gene as an editing target because it has been ever taken for evaluating the effect of donor sizes on genome editing efficiency in our previous work, where good performance was got with one of the tested plasmids, pKO-*ZMO0038*-3 carrying a 600-bp donor DNA (Zheng et al. 2019). We thus constructed the editing plasmids based on pKO-*ZMO0038*-3. Since paired crRNAs simultaneously targeting two genomic loci were required for double nicking, a new editing plasmid, pKO-*ZMO0038*n, was constructed to bear an artificial CRISPR array consisting of two spacers derived from different strands and three insulating direct repeats. Two different crRNA guides were to be produced from the plasmid-borne artificial CRISPR and were expected to direct a pair of Cascade-nCas3 units to introduce double nicks on different strands of the target, generating a DSB with an overhang (**Fig. 2a**).

**Fig. 2.**
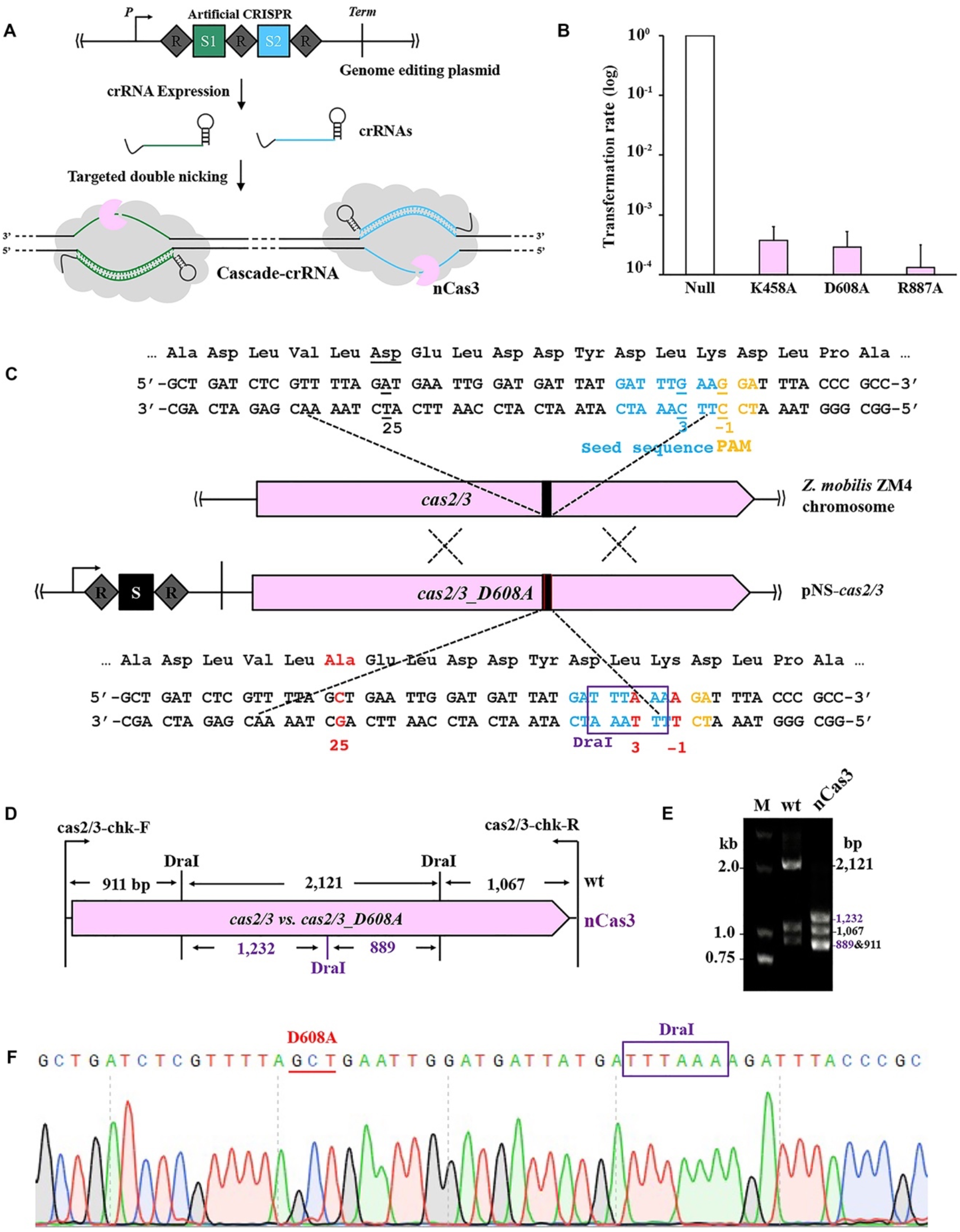
Establishment of a Cascade-nCas3-mediated genome editing tool. (**A**) A genome editing plasmid contained an artificial CRISPR locus consisting of two spacers (S1 and S2) and three insulating direct repeats (R). Paired self-targeting crRNAs were to be produced from the artificial CRISPR and simultaneously guide Cascade complexes to bind to two target sequences matching S1 and S2, respectively, located on opposing strands. The nCas3s were then recruited to nick the dsDNA within the target sequences. (**B**) Transforming competent cells of the Δ*cas2/3* strain with *ZMO0038* knockout plasmids each expressing a Cas3 nickase mutant (K458A, D608A, or R887A). Transformation rates are present as relative values to that with a reference plasmid with no Cas3-encoding gene (Null), the latter of which was set to be 1.0. Experiments were performed in triplicate. Error bars represent the standard deviation of the mean. (**C**) Schematic showing nucleotide substitution of *cas2/3*. The spacer in the genome editing plasmid (pNS-*cas2/3*) for nucleotide substitution of *cas2/3* and the corresponding protospacer in *cas2/3* are indicated as a black box. The PAM motifs are shown in orange while the seed sequence in crane. The designed mutations are indicated as red fonts in *cas2/3_D608A*, whereas the corresponding original nucleotides are underlined in *cas2/3.* The restriction site for DraI (TTTAAA) that is to be introduced is framed in a purple box. (**D**) Schematic showing the digestion sites by Dra, among which the newly introduced one is in purple, in the PCR fragments amplified by a primer set of cas2/3-chk-F and cas2/3-chk-R. the predicted sizes of digestion products are indicated. (**E**) Electrophoretic analysis of DraI-treated colony PCR products amplified from the wild-type strain (wt) and the mutant candidate (nCas3) using primers shown in (**D**). M, DNA size marker. (**F**) Representative chromatographs of Sanger sequencing confirming the designed nucleotide substitutions in *cas2/3.*

Initially, taking the convenience of protein expression via an episomal vector, we cloned each gene encoding an nCas3 variant to pKO-*ZMO0038*n, yielding three editing plasmids, pKO-*ZMO0038*-K458A, pKO-*ZMO0038*-D608A, and pKO-*ZMO0038*-R887A (**Table S1**). These editing plasmids, and the cloning vector pEZ15Asp as a reference (Yang et al. 2016), were then individually electroporated into *Z. mobilis* Δ*cas2/3*, a previously constructed *cas2/3* knockout (Zheng et al. 2019). Only very few transformants could be yielded from transformations with the editing plasmids, showing hundreds-fold lower transformation rates than that with the reference plasmid (**Fig. 2b**) and thereby reflecting a potent killing effect of the nCas3s on the host cells.

Speculatively, overexpression of the nCas3 variants was toxic to *Z. mobilis* cells. To verify this speculation, we removed the artificial CRISPR from the editing plasmids, generating three expression plasmids, pEZ-K458A, pEZ-D608A, and pEZ-R887A (**Table S1**), with each expressing a corresponding nCas3 whereas no crRNA production. We failed in yielding any transformant from the transformations with these expression plasmids (**Table 1**), suggestive of strong cytotoxicity of the nCas3s *per se* to the *Z. mobilis* cells.

**Table 1.** Transformation rates (TR) and editing efficiencies (EE) of various genome-editing plasmids in *Z. mobilis* DRM1 and DRM2, respectively.

| Plasmid | TR (cfu/μg DNA) |  | EE [% (editing/tested)] |  |
| --- | --- | --- | --- | --- |
|  | DRM1 | DRM2 | DRM1 | DRM2 |
| pEZ15Asp | $(3.21 \pm 1.53) \times 10^6$ | $(2.33 \pm 1.23) \times 10^6$ | - | - |
| pKO-ZMO0038n | - | $(4.09 \pm 1.14) \times 10^5$ | - | 100 (16/16) |
| pKO-ZMO0252 | $(9.49 \pm 0.51) \times 10^2$ | $(2.47 \pm 0.65) \times 10^5$ | 37.5 (6/16) | 93.75 (15/16) |
| pDel-10k | $(1.51 \pm 0.51) \times 10^3$ | $(3.02 \pm 0.83) \times 10^4$ | 31.25 (5/16) | 87.5 (14/16) |
| pRMV | - | $(7.26 \pm 0.25) \times 10^4$ | - | 93.75 (15/16) |

- : Not determined

Indeed, it was reported that, if not properly controlled, endonucleases in CRISPR-Cas systems provided protection with the risk of toxic activity against the host (Leon et al. 2018). Bacteria have therefore evolved different mechanisms to modulate the activity of Cas nucleases. For example, in Type I-F systems four Cas1 molecules form a complex with two molecules of Cas2-Cas3 fusion to neutralize the nuclease activity of the latter (Rollins et al. 2017). Reasonably, such a balance might be broken by the overproduction of a Cas3 nickase that disrupted the certain ratio between the subunits.

### A CRISPR-nCas3 genome editing tool is established upon an *in situ* nCas3 variant

In order to attain genome editing with the CRISPR-nCas3 system, we next sought to generate an *in situ* nCas3 by introducing alanine substitution of the D608 residue. To this end, a genome editing plasmid, pNS-*cas2/3* for nucleotide substitutions of *cas2/3*, was designed. By carefully inspecting the coding sequences in the vicinity of the D608 residue, a 5’-TCC-3’ PAM located on the template strand was found and therefore the 32-nt sequence immediately downstream of it was considered as a protospacer (**Fig. 2c**).

Three nucleotide changes were introduced into the donor DNA, that is, C-1T, C3T, and T25G, where for clarity, we defined the numbering scheme for protospacer positions as following: the position immediately downstream of PAM is called 1, with subsequent positions being 2, 3, *etc.*, up to 32; while positions within the PAM are referred to as −1, −2, and −3, with −1 is the closest to the protospacer. The C-1T and C3T substitutions interrupted the functional 5’-TCC-3’ PAM and the seed sequence to allow for cell surviving after editing, which did not result in any change of protein sequences; whilst the T25G mutation resulted in altering the original GAT codon for aspartic acid (D) to the GCT codon for alanine (A). In addition, the C3T mutation led to the formation of a TTTAAA restriction site for the DraI endonuclease (**Fig. 2c**). This allowed us to rapidly screen strains with expected edits by colony PCR amplification of DNA fragments encompassing the edited region followed by DraI treatment of the PCR products.

More than 200 transformants were yielded after transforming the pNS-*cas2/3* plasmid into the DRM1 cells (Zheng et al. 2019). Using the primer set of *cas2/3*-chk-F and *cas2/3*-chk-R (**Supplementary Table S2**), DNA fragments of 4,099 bp were amplified from 4 randomly picked transformants. The PCR products were then digested with DraI followed by electrophoretic analysis using an agarose gel. DraI treatment of the reference sample would produce 3 bands with the sizes of 911 bp, 2,121 bp, and 1,067 bp, respectively. If the modifications correctly occurred, an additional DraI restriction site would be introduced in the 2,121-bp fragment, such that the 2,121-bp DNA would be further cut into two fragments of 1,232 bp and 889 bp by DraI (**Fig. 2d**). The results suggested that the designed *in situ* nCas3 was successfully generated and confirmed via analyses of DraI treatment and Sanger sequencing of the PCR products (**Fig. 2e,f**).

The resulting Cas3(D608A) strain, designated *Z. mobilis* DRM2, was then used as the genetic host for CRISPR-nCas3 genome editing. Knockout of *ZMO0038* was attempted in *Z. mobilis* DRM2 cells to assess the capability of CRISPR-nCas3 in genome editing. Transformation of DRM2 competent cells with the pKO-*ZMO0038*n yielded hundreds of transformants, showing a transformation rate of only about 10-fold lower than that with the reference plasmid (**Table 1**). As expected, after HDR of the DSB generated through double nicking by a pair of Cascade-nCas3 units, deletion of the target gene would occur (**Fig. 3a**). Of the obtained transformants, 16 were randomly picked up and analysed by colony PCR and Sanger sequencing genotypic characterization. The results showed that all the tested transformants were identified to harbour the designed deletion of *ZMO0038* (**Fig. 3b,c**), giving an editing efficiency of 100% (**Table 1**).

**Fig. 3.**
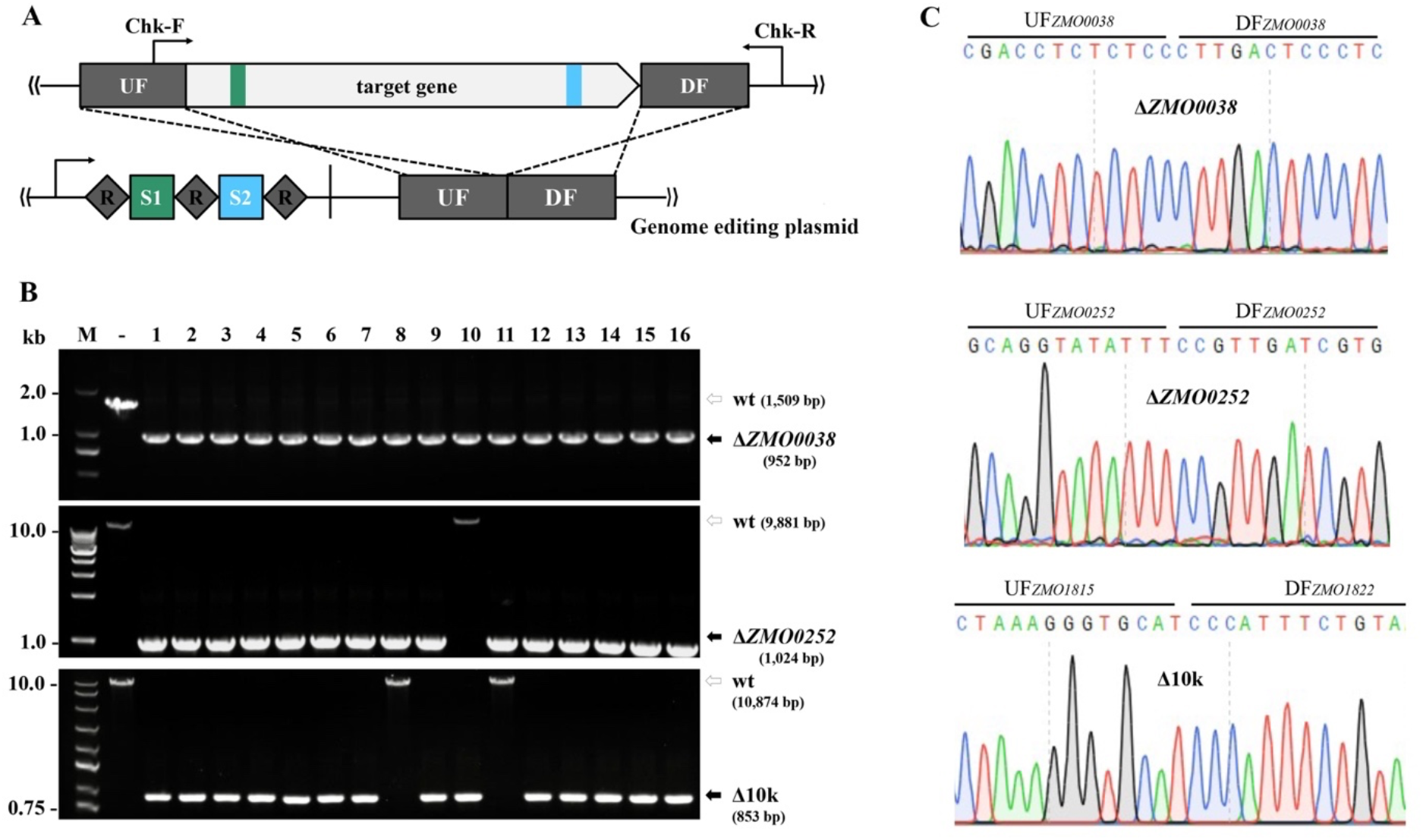
Efficient genome editing using CRISPR-nCas3. (**A**) Schematic showing design of the genome editing plasmid. An artificial CRISPR expressing two targeting crRNAs and a donor DNA consisting of an up-flanking (UF) and a down-flanking (DF) DNA stretches of the target gene are contained in the all-in-one editing plasmid. (**B**) Colony PCR screening of deletion mutants of *ZMO0038* (upper panel), ZMO0252 (middle), a ~10-kb genomic fragment (lower panel), respectively, using a gene-specific primer set, Chk-F/ChkR, as indicated in (**A**). Predicted sizes of PCR products in wild-type (wt) and the expected deletion mutants (Δ*ZMO0038*, Δ*ZMO0252* or Δ10k) are indicated with unfilled and filled black arrows, respectively. -, PCR amplification using genomic DNA of *Z. mobilis* ZM4 as a DNA template; M, DNA size marker. (**C**) Representative chromatographs of Sanger sequencing results.

We noticed that transformation of DRM2 with pKO-*ZMO0038*n got a rate of about 10-fold higher than that obtained from transformation of DRM1 cells with the pKO-*ZMO0038* plasmid in our previous study (Zheng et al. 2019). Although in both cases the efficiencies of *ZMO0038* knockout were of 100%, the latter was attained by improving pKO-*ZMO0038* transformation rate through destroying a restriction-modification (R-M) system (Zheng et al. 2019). The further enhanced pKO-*ZMO0038*n transformation rate might reflect a greater capability of the CRISPR-nCas3 in genome editing. To corroborate this, we constructed the pKO-*ZMO0252* plasmid by taking the same strategy as illustrated in **Fig. 3a** to delete the 8,955-bp *ZMO0252* gene encoding a component of a predicted Type I secretion system (Zhang et al. 2019), looking at whether the CRISPR-nCas3 could also mediate efficient removal of larger genomic fragments. Transforming pKO-*ZMO0252* into DRM2 cells yielded hundreds of transformants. Among them, 16 were randomly chosen and 15 out of which were identified to be edited cells with the desired genotypes (**Fig. 3b,c**), showing an editing efficiency of 93.75% (15/16) (**Table 1**). Strikingly, an efficiency of 87.5% was also yielded in the experiment of deleting the 10,021-bp genomic fragment that we took as an editing target in our previous work (Zheng et al. 2019) (**Fig. 3b**; **Table 1**). We also used these editing plasmids to perform the same genome editing options in DRM1 cells using the CRISPR-Cas3 tool, yielding editing efficiencies of 31.25% and 37.5% for deletion of *ZMO0252* and 10-kb fragment, respectively. Particularly, for the 10-kb fragment deletion experiment, both the transformation rates of editing plasmid and the editing efficiency are comparable to that seen in our previous study (**Table 1**). These results demonstrated the overall reproducibility of the observed high-efficiency editing via CRISPR-nCas3.

### CRISPR-nCas3 enables simultaneous removal of large genomic fragments

To further illustrate the versatility of this CRISPR-nCas3-based technology, we opted to use it for simultaneously removing two large genomic loci using a single editing plasmid, pRMV (**Fig. 4a**). After electroporating pRMV into DRM2 cells, hundreds of transformants appeared on the selective plate, getting an average transformation rate of (7.26±0.25) × 10^4^ CFU/μg plasmid DNA (**Table 1**). Of the obtained transformants, 16 were randomly selected for genotypic characterization by colony PCR analysis using specific primer sets listed in **Table S2**. As shown in **Fig. 4b**, 13 colonies (*i.e*. Strains 1-5, 7-9, 11, and 13-16) contain the 10-kb fragment deletion, while 14 colonies (*i.e.* Strains 2-9 and 11-16) are *ZMO0052* knockouts. Collectively, a total of 15 colonies carry at least one deletion, giving an overall editing efficiency of 93.75%. Notably, 12 strains contain both the deletions, showing an engineering efficiency of 75% (**Fig. 4c**).

**Fig. 4.**
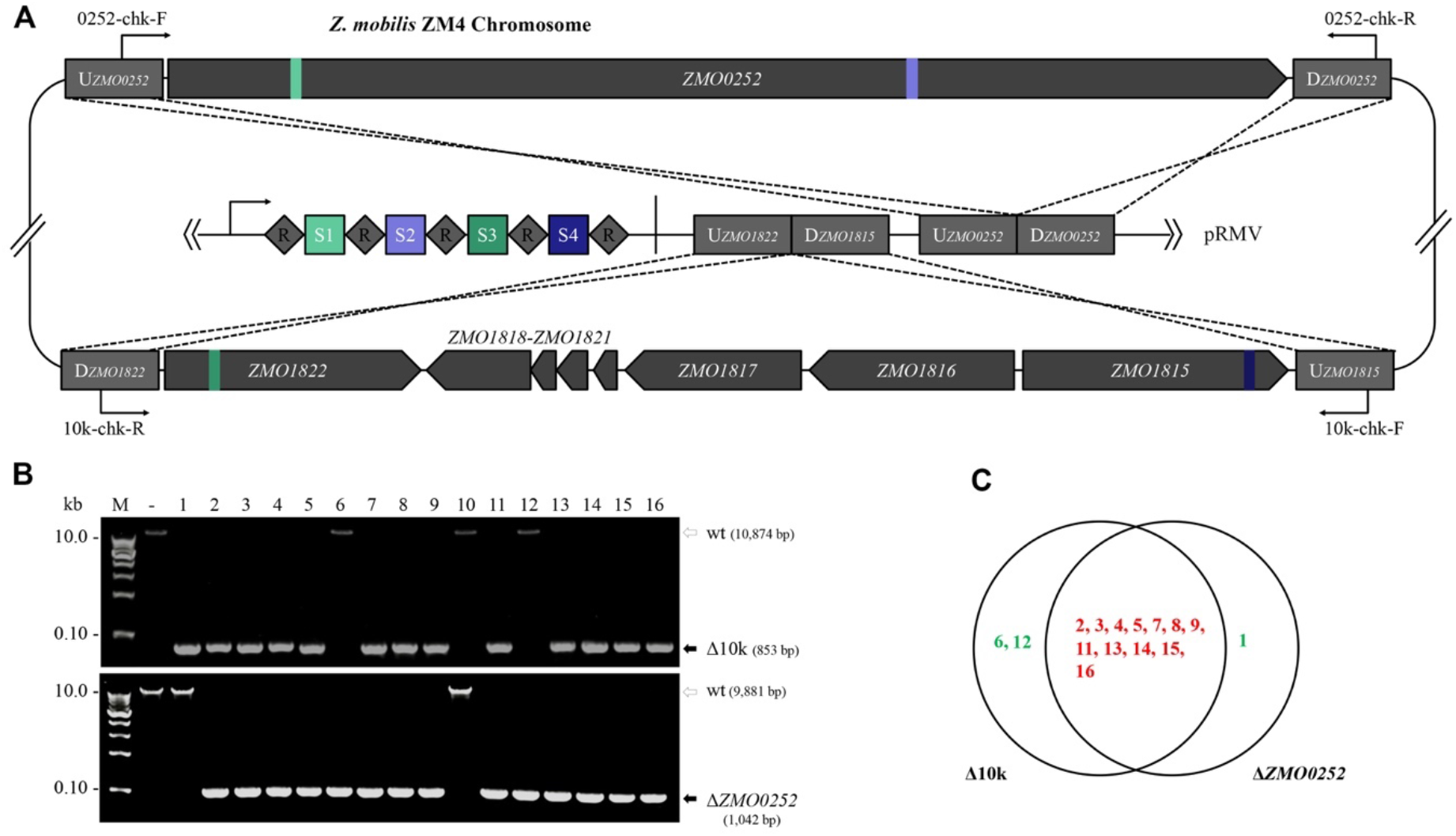
Simultaneous removal of two large genomic fragments using CRISPR-nCas3. (**A**) Schematic showing design of an 8,995-bp ZMO0052 gene and a ~10-kb genomic fragment (spanning genes of *ZMO1815-ZMO1822*) deletion. The pRMV encodes four spacers with S1 and S2 matching sequences within the *ZMO0052* gen while S3 and S4 within the 10-kb region, respectively. DNAs up-flanking (UF) and down-flanking (DF) of the targets were concatenated on the same plasmid as recombination donors. (**B**) Colony PCR screening of deletion mutants of the10k genomic fragment (upper panel) and *ZMO0052* (lower panel), respectively, using specific primer sets as indicated in (**A**). Predicted sizes of PCR products in wild-type (wt) and the expected deletion mutants (Δ10k or Δ*ZMO0252*) are indicated with unfilled and filled black arrows, respectively. -, PCR amplification using genomic DNA of *Z. mobilis* ZM4 as a DNA template; M, DNA size marker. (**C**) Distribution of genomic deletions in the tested transformants. Transformants with both deletions or single deletion are shown in red and green fonts, respectively.

## Discussion

This work reports the first establishment, to the best of our knowledge, of an advanced CRISPR-nCas3 genome editing method in *Z. mobilis*, which includes a Cas3 nickase. Differently from the Cas9 nucleases which use two nuclease domains, an NHN and a RucV, to respectively cleave the different strands of a dsDNA target (Cong et al. 2013), Cas3 proteins in Type I systems use only one ssDNA nuclease domain to gradually nick the two strands(Sinkunas et al. 2011). As previously demonstrated, Cas3 is recruited to a target upon formation of an ssDNA-containing R-loop through crRNA-directed Cascade-binding and cuts the displaced ssDNA strand first; whilst cleavage of the crRNA-paired strand requires its ATP-dependent helicase domain to unwind the dsDNA target (Sinkunas et al. 2011). This feature allows us to generate the Cas3 nickase mutants by inactivating the helicase domain of the Cas3 nuclease-helicase. Interestingly, as there are several residues essential for the helicase activity (Sinkunas et al. 2011; Gong et al. 2014), it is flexible to create different nickase mutants by inactivating any of the essential residues. By contrast, an nCas9 can only be a mutant of either a D10A in RuvC or a H840A in HNH (Ran et al. 2013). As derived from an endogenous system, it is more convenient to simultaneously produce crRNA pairs, which is an important requirement for nCas-mediated genome editing (Ran et al. 2013), through processing the precursor RNAs of the single artificial CRIPSR by the Csy4/Cas6f protein (Przybilski et al. 2011).

Given the fact that enhanced DNA targeting specificity was achieved with a CRISPR-nCas9 (Ran et al. 2013), the same should be also true for this CRISPR-nCas3, being of increased genome editing specificity. Also, as the nCas9 showed an obvious advantage in helping base editing over other Cas9 variants (Nishida et al. 2016), we envision that nCas3-based toolkits, such as base editors, would be soon available for various bacteria harbouring an active Type I CRISRP-Cas. Furthermore, very recently a Type I-C system has been evidenced for genome editing in several bacteria(Csorgo et al. 2020). Type I-F systems have relatively fewer Cas components among the Type I subtypes (Makarova et al. 2020), they thereby could be also readily potable for heterologous genome editing in other organisms.

Significantly elevated editing efficiencies (near-100%) were observed in the application of CRISPR-nCas3 tool for genome editing including simultaneous deletion of large genomic fragments. Our previous demonstrations showed that only up to 50% efficiency for removal of one large genome fragment could be attained, and simultaneous deleting multiple small DNA stretches yielded an efficiency of 18.75%. We noticed that, for simultaneous removal two large genome fragments, the transformation rate of the editing plasmid and the engineering efficiency are at the same level as that observed for deletion of either of them, indicating that simultaneously deleting more genomic targets would be also efficiently achieved with this CRISPR-nCas3 tool. Since editing efficiencies rely largely on the repair rates of DSBs by the host’s repair systems, together with the fact that *Z. mobilis* lacks an NHEJ system, the enhancement of editing efficiency might be due to faster repair of the DSBs by the HDR systems, thereby letting more cells be recovered from self-targeting. Possibly, the DSB ends produced by nCas3-mediated double nicking each carries an overhang structure, which might be more efficiently sensed and bound by RecA to initial DSB repair (Wigley 2013). Another possibility could be also that the overhangs might trigger or activate an alternative repair system with an even higher efficacy, as bacteria generally possess multiple HDR systems (Bernheim et al. 2019), for instance *Z. mobilis* ZM4 encodes at least two HDR mechanisms, *i.e.* an AddAB and a RecF (Yang et al. 2018). By the way, this work offers an easy method to produce DSBs at defined genomic locations with expected terminal structures for studying HDR mechanisms in bacteria *in vivo*. Other possibilities include that double nicking by nCas3 might be lesser toxic than processive degradation by Cas3 nuclease-helicase, thus enabling more cells to be recovered. Bacteria are generally sensitive to CRISPR-mediated chromosomal self-targeting. Potent CRISPR self-targeting may lead to failure in yielding any recovered cells with the designed edits. Indeed, in this work, transformation of the same editing plasmid into cells with an nCas3 background yielded about 20-fold higher rate than into those with a Cas3 background (**Table 1**). In the future, comprehensive studies, combining structural, genetic and biochemical analyses, on the HDR mechanisms in *Z. mobilis* may offer molecular explanations for the observed phenomenon, as well as mechanistic insights for directing high-efficiency genome editing.

Conclusively, we have created a Type I-F CRISPR-nCas3-based technology that represents currently the most efficient and straightforward genome engineering tool for the important industrial bacterium *Z. mobilis*. It has allowed us to achieve highly efficient removal of genomic fragments in a large-scale manner in *Z. mobilis*, and hence would expedite the development and improvement of this bacterium as an ideal chassis for synthetic biology researches. This study expands the available tools for CRISPR-mediated genome engineering and may serve as a framework for future development of next-generation CRISPR-Cas technologies.

## Methods

### Strains, growth conditions and electroporation of *Z. mobilis*

*Z. mobilis* ZM4 and derivatives constructed in this work were listed in **Supplementary Table S1**. *Z. mobilis* strains were grown at 30°C in an RMG medium (20 g/L glucose, 10 g/L yeast extract, 2 g/L KH_2_PO_4_). If required, spectinomycin was supplemented to a final concentration of 200 μg/mL for *Z. mobilis* and 50 μg/mL for *Escherichia coli*. Competent cells of *Z. mobilis* were prepared as previously described (Yang et al. 2016) and transformed with plasmids by electroporation using Bio-Rad Gene Pulser (0.1-cm gap cuvettes, 1.6 kV, 200 Ω, 25 μF) (Bio-Rad, Hercules, CA, USA) following the method developed for *Z. mobilis* (Okamoto and Nakamura 1992). Electroporated cells were incubated in an RMG2 medium for 3 hours at 30°C prior to plating.

### Construction of plasmids

Artificial CRISPR expression plasmids were constructed based on the *E. coli-Z. mobilis* shuttle vector, pEZ15Asp (Yang et al. 2016). A DNA block consisting of the leader sequence of the chromosomal CRISPR2 as a promoter and three CRISPR repeats separated by two BsaI and two BsmBI restriction sequences in opposite orientation, respectively, was synthesized from GenScript (Nanjing, China) and used as a template for PCR amplification with the primer pair of L3R-XbaI-F/L3R-EcoRI-R. Then, the PCR product was digested with XmaI and BamHI and subsequently inserted into the pEZ15Asp vector, generating the base vector pL3R. Digestion of pL3R with BsaI generated a linearized plasmid having protruding repeat sequences of 4 nt at both ends. Double-stranded spacer DNAs were prepared by annealing two spacer oligonucleotides through being heated to 95°C for 5 min followed by cooling down gradually to room temperature. Likewise, the second spacer could be inserted in between of the repeats by using the BsmBI sites. The spacer fragments were designed to correspondingly carry protruding ends complementary to those in the linearized vector. Therefore, self-targeting plasmids each bearing an artificial CRISPR with two self-targeting spacers were generated by gradually ligating spacer inserts with the linearized vectors. By repeating the reactions, the pRMV plasmid for simultaneous remove of the two large genomic fragments was yielded. Subsequently, donor DNA fragments each containing a mutant allele of a target gene were generated by splicing and overlap extension PCR (SOE-PCR) (Horton et al. 1990) and individually cloned into their cognate self-targeting plasmids through the T5 exonuclease-dependent DNA assembly (TEDA) method (Xia et al. 2019). Genome editing plasmids for creating the nCas3 mutants were constructed based on the pL2R plasmid vector following the previously described method (Zheng et al. 2019).

Expression plasmids of Cas3 and Cascade proteins were constructed with the *E. coli* pET28a expression vector. Individual *cas* gene was PCR-amplified from *Z. mobilis* total DNA using specific primers listed in **Table S1**. The PCR product of *cas3* gene was used as a template to amplify the mutant genes through splicing and overlap extension PCR (SOE-PCR) (Horton et al. 1990) using primers listed in **Table S2** containing the corresponding mutations. After digested with the enzymes indicated in each PCR primer, the DNA fragments were individually cloned to pET28a at compatible sites, giving pET-Cas3, pET-Cas5, pET-Cas6, pET-Cas7, pET-Cas8, pET-K458A, pET-D608A, and pET-R887A.

For overexpression of the Cas3 variants in *Z. mobilis*, each gene was amplified from the pET-K458A, pET-D608A, and pET-R887A, respectively, using specific primers listed in **Table S2**, and clone to the pEZ15Asp vector or the genome editing plasmid pKO-*ZMO0038*n at EcoRI and XbaI sites, yielding pEZ-K458A, pEZ-D608A, and pEZ-R887A, or pKO-*ZMO0038*-K458A, pKO-*ZMO0038*-D608A, and pKO-*ZMO0038*-R887A, respectively.

All plasmids were listed in **Table S1**. All oligonucleotides were synthesized from GenScript (Nanjing, China) and listed in **Table S2.** Restriction enzymes and T5 exonuclease were purchased from New England Biolabs (Beijing) Ltd (Beijing, China).

### Expression and purification of Cas proteins

The Cas expression plasmids were individually transformed into *E. coli* BL21 (DE3) and expression of the His-tagged Cas proteins was performed following the instruction of the protein purification kit (Qiagen, Valencia, CA, USA). Single colonies of transformed cells were cultivated overnight, followed by 1/100 dilution into 100 mL of LB media containing 100 μg/mL ampicillin. The cells were firstly incubated at 37°C to an OD_600_ of 0.6-0.8, then transferred to a shaker and induced with isopropyl β-D-1-thiogalactopyranoside (IPTG) in a final concentration of 0.5 mM at 16°C. After continuously shaking for 22 hours, cells were harvested, lysed, and purified using Ni-NTA resin (Qiagen, Valencia, CA, USA). The purified proteins were desalted with desalting column (GE Healthcare, Chicago, IL, USA) using AKTA system (GE Healthcare, Chicago, IL, USA), and finally confirmed by SDS-PAGE electrophoresis.

### Plasmid DNA cleavage assay

One hundred and fifty ng of the pL2R plasmid DNA was incubated at 30°C with 250 nM of Cas3 or one of the nCas3 variants, a crRNA carrying a spacer targeting a 5’-CCC-3’ PAM-preceded 32-nt sequence of pL2R, and the Cascade proteins in a reaction buffer containing 2 mM MgCl_2_ and 0.5 mM ATP. The reaction products were checked by agarose gel electrophoresis. The crRNA was synthesized from GenScript (Nanjing, China) and listed in **Table S2**.

### Construction and screening of mutants, and curing of genome editing plasmids

The genome editing plasmids were individually introduced into *Z. mobilis* cells through electroporation. Electroporated cells were spread on RMG agar plates containing spectinomycin at a final concentration of 200 μg/mL (RMGSp) and incubated at 30°C until colonies were observed. Mutant candidates were screened by colony PCR using primers listed in **Table S2**. The resulting PCR products were analysed by agarose gel electrophoresis and confirmed by Sanger sequencing (GenScript, Nanjing, China). The genome editing plasmids were cured following the method we previously developed (Zheng et al. 2019).

## Availability of data and materials

The authors declare that the main data supporting the findings of this work are available within the article and its supplementary information files or from the **corresponding** authors upon reasonable request.

## Authors’ contributions

WP, YZ, SY, and LM designed the research; YH, QW, and JL performed the experiments; WP, YH, and YZ wrote the manuscript. All authors contributed to data analyses, read, revised and approved the final manuscript.

## Acknowledgements

This work was supported by the Scientific Research Program of Hubei Provincial Department of Education (Q20161007), the National Key Technology Research, the Development Program of China (2018YFA0900300), the National natural Science Foundation of China (U1932141), the Hubei Technical Innovation Special Fund (2019AHB055 and 2018ACA149), and the Innovation Base for Introducing Talents of Discipline of Hubei Province (2019BJH021). WP acknowledges the support from the State Key Laboratory of Biocatalysis and Enzyme Engineering.

